# Adaptation by stochastic tuning of gene expression in mammalian cells

**DOI:** 10.1101/2025.05.07.651953

**Authors:** Amir Momen-Roknabadi, Panos Oikonomou, Saeed Tavazoie

## Abstract

Cells often face environmental challenges for which they lack pre-programmed regulatory responses. We have previously shown that yeast cells can, nevertheless, adapt to such unfamiliar conditions by a phenomenon we have termed stochastic tuning. In this process, cells use gene expression noise to randomly change the expression of individual genes and actively reinforce those changes that improve the overall health of the cell. Here, we provide experimental evidence for stochastic tuning in human cells adapting to the lethal chemotherapeutic agent methotrexate. As in yeast, the tuning process is not driven by mutations and is reversible upon removal of methotrexate challenge. We show that stochastic tuning is a conserved eukaryotic mechanism of cellular adaptation, and potentially a key factor in the phenomenon of cancer cell plasticity underlying chemotherapy resistance.

## Introduction

Cells adapt to changes in their external environment primarily by adjusting the expression of their genes. Our modern understanding of cellular adaptation goes back to the seminal work of Jacob and Monod^1,2^. They were the first to describe that a genetically encoded regulatory program activates the expression of the Lac operon in the presence of lactose and absence of the preferred sugar glucose^2^. The Jacob and Monod framework has led directly to our modern conception of gene regulatory networks (GRNs)^3^. Evolved over geological timescales, GRNs perceive recurring patterns of stimuli in the native habitat and activate dedicated gene regulatory programs. However, cells often encounter novel extreme environments for which predefined regulatory programs do not exist. How do cells, nevertheless, adapt to such unfamiliar contingencies? A clinically relevant example is the ability of cancer cells to adapt to inhibitory/lethal xenobiotic agents. Resistance to such chemotherapy drugs often occurs through global modulation of select genes—resistance inducing genes are upregulated and those that make the cells sensitive are downregulated^4-11^. Although DNA mutations are a major source of such adaptation, there is increasing evidence for non-mutational processes in chemotherapy resistance^10-17^. The underlying mechanisms of such non-genetic adaptation phenomena are not well understood. More generally, a major emerging question is whether there exist novel mechanisms by which cells can adapt to extreme/unfamiliar environments where their native sensory and regulatory systems are no longer adequate.

Our laboratory has recently described a novel regulatory mechanism in *Saccharomyces cerevisiae*, which allows cells to adapt to unfamiliar environments beyond the capacity of their genetically encoded regulatory programs. In this phenomenon, called stochastic tuning, it is proposed that cells utilize transcriptional noise to randomly change the expression of individual genes, and reinforce those changes that improve the overall health of the cell^18^ (**Figure 1**). Importantly, this process is distinct from population-level selection underlying stochastic phenomena such as bacterial persistence^19,20^. Stochastic tuning may have originally evolved as a mechanism for adaptation of single-cell organism to extreme environments. However, it may have later found utility in multicellular organisms, optimizing cell states to facilitate physiological homeostasis under conditions of extreme challenge.

**Figure 1.**
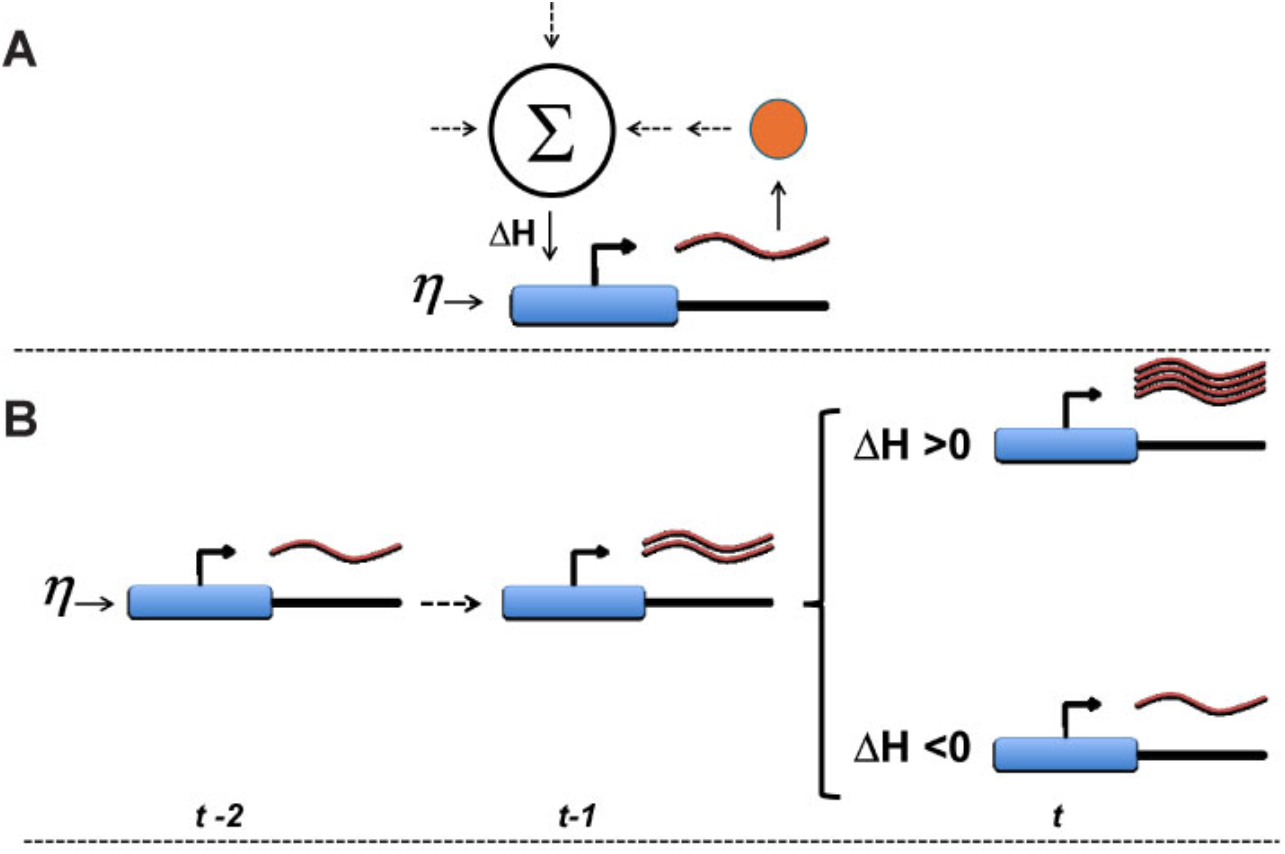
Stochastic tuning: cellular adaptation by noise-driven optimization of cell-health. **(A)** Working model of stochastic tuning. Individual genes can exhibit random bursts of transcription (*η*), a temporary record of which is retained in the modification state of local chromatin. A proposed global health integrator (Σ) continuously broadcasts a signal (ΔH) to all promoters, conveying whether the overall health of the cell is improving or deteriorating. (**B**) A simple example of stochastic tuning can be seen for a gene experiencing a random burst of transcription. If this is followed by an improvement in overall health (ΔH>0), the expression apparatus further increases transcriptional output. Conversely, if health deteriorates (ΔH<0), the expression apparatus decreases transcriptional output. This process leads to establishment of expression levels that maximize the overall health of the cell. Adapted from (Freddolino et al. *eLife* (2018) doi: 10.7554/eLife.31867)

Resistance of cancer cells to xenobiotic chemotherapy agents is an example of adaptation to extreme/unfamiliar environments that may be beyond the capacity of genetically encoded regulatory programs. Generally, it is thought that resistance is due to pre-existing or acquired mutations^21^. However, in many cases, cells become resistant by modulating the levels of the drug’s target or resistance-inducing factors through a non-mutational process of unknown mechanism. For example, ATP-binding cassette (ABC) transporter family of transmembrane proteins control the flux of many biological substrates. By increasing the expression levels of the members of this family, cells can become resistant to multiple chemotherapy agents simultaneously^8,21-23^. However, the mechanism by which cells specifically adjust the expression level of relevant resistant genes is not currently understood. Franca et al.^11^ observe that this form of non-genetic resistance is accompanied by a progressive increase in cell fitness along a ‘resistance continuum’. Furthermore, Li et al.^10^ show that epigenetic therapy resistance can induce long-term maintenance of allele specific gene expression concomitant with increasing cell fitness. These phenomena strongly suggest that stochastic tuning may be the key underlying mechanism of epigenetic cancer therapy resistance. To investigate stochastic tuning in mammalian cells, we engineered a chromosomally integrated reporter system based on resistance to methotrexate (MTX), an antimetabolite drug that inhibits dihydrofolate reductase (DHFR), a critical enzyme in purine biosynthesis^24,25^. We show that a mechanism consistent with stochastic tuning enables human HEK293 cells to upregulate a resistant DHFR variant and grow under otherwise lethal levels of MTX.

## Results

### A mammalian model of stochastic tuning

To study the role of stochastic tuning in the adaptation of mammalian cells to extreme environmental challenges, we developed a model system using human HEK293 cell lines (Flp-In293 variant, Thermofisher) and their response to MTX (**Figure 2A**). Due to their vast available genetic tools, HEK293 cells have been widely used as a model to study many aspects of molecular biology. MTX is an inhibitor of dihydrofolate reductase (DHFR). DHFR converts dihydrofolate to tetrahydrofolate, a key step in purine biosynthesis^24^. MTX has been widely used for treatment of various cancers. However, its use has been limited due to emergence of resistance. In some cases, resistance is due to mutations in DHFR that reduces the binding of MTX^24-26^. We used one such variant, the murine DHFR (mDHFR) with Leu22Arg mutation, rendering it resistant to MTX^27^. We created a construct based on pCDNA5/FRT (Thermofisher) where a fusion of mDHFR with mRuby fluorescent protein is placed under the control of a weak synthetic promoter (tetracycline (tet)-off *cis-*regulatory system and a minimal CMV promoter) and transfected into Flp-In293 cells to select for chromosomally integrated reporters. Here, we will refer to the line with mDHFR-mRuby in the absence of the inducer tTA as the “reporter line”. The tet-off system has a very low level of expression, since our cell line does not contain the tTA (tet activator). In addition, we generated a “positive control” line, containing the mDHFR-mRuby reporter, and constitutively expressing tTA (“positive control”). As a “negative control”, we engineered a line that expresses a non-functional form of mDHFR-mRuby with an in-frame deletion of DHFR active site (**Figure 2B**).

**Figure 2.**
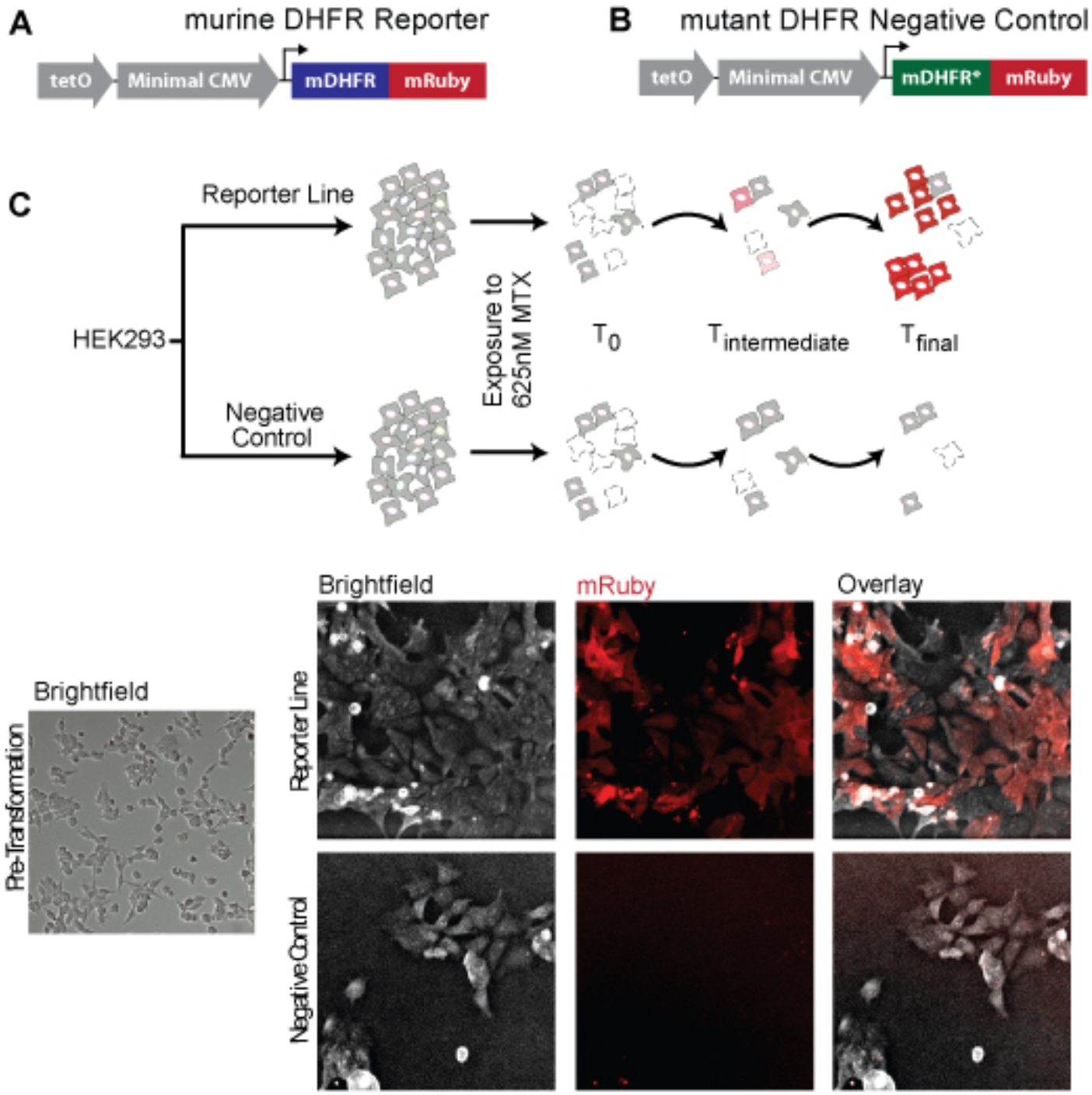
Mammalian cells exhibit stochastic tuning. (**A**) mDHFR-mRuby fusion is cloned downstream of a tet operator (tetO) region and a minimal CMV promoter. (**B**) The negative control construct has a mutated mDHFR-mRuby fusion under the identical synthetic promoter. (**C**) FLP-In293 cells were transformed with either the mDHFR-mRuby or non-functional form of mDHFR-mRuby (negative control). Following exposure to MTX, the cells with mDHFR-mRuby tune the expression of mDHFR-mRuby to high levels while the negative control cells do not. The micrographs are adjusted for brightness and contrast.

### HEK293 model shows stochastic tuning

As the first step, we determined the proper concentration of MTX that is lethal but allows a fraction of cells to survive for long periods of time. We exposed all three of the above lines to different concentrations of MTX (0-10 μM) and measured the cells’ survival over the span of a week. We found that at the concentration of 625 nM, ∼95% of cells with only the mDHFR or nonfunctional-mDHFR transgene, or with both nonfunctional -mDHFR and tTA transgenes died. These lines, however, had some microcolonies, exhibiting minimal growth. In the case of the line with both functional mDHFR and tTA transgenes (positive control), the cells grew almost without any cell death at 625 nM of MTX. In addition, this positive control cell line exhibited high mRuby expression for an extended period. Remarkably, following 15-18 days of drug exposure, we observed a fraction (∼1%) of colonies in the reporter line that grew much faster than the other surviving microcolonies and were positive for mRuby expression. We did not observe any colonies that were growing in the negative control line at 625 nM of MTX (**Figure 2C**).

Since there could be genetic heterogeneity between the tuned colonies, we isolated individual surviving colonies from the reporter line in MTX. These cells divide approximately once every 24 hours at 625 nM of MTX. Flow cytometry analysis showed a substantial increase (>100-fold) in the expression of the mDHFR-mRuby (**Figure 3A**). MTX acts as a competitive inhibitor of DHFR^28^. Therefore, we hypothesized that by exposing cells already tuned to 625 nM of MTX, to further increasing concentration of MTX, we may observe corresponding increases in mDHFR-mRuby expression in tuned cells. To test this, we exposed the tuned cells to 1.25 μM, 2.5 μM or 5 μM of MTX. Next, using flow cytometry, we measured the levels of mRuby expression over 24 days. As can be seen in **Figure 3A**, after 24 days of exposure to higher concentrations of MTX, tuned cells showed statistically significant increases in mDHFR-mRuby expression (*p*-value<0.001, Mann-Whitney U Test).

**Figure 3.**
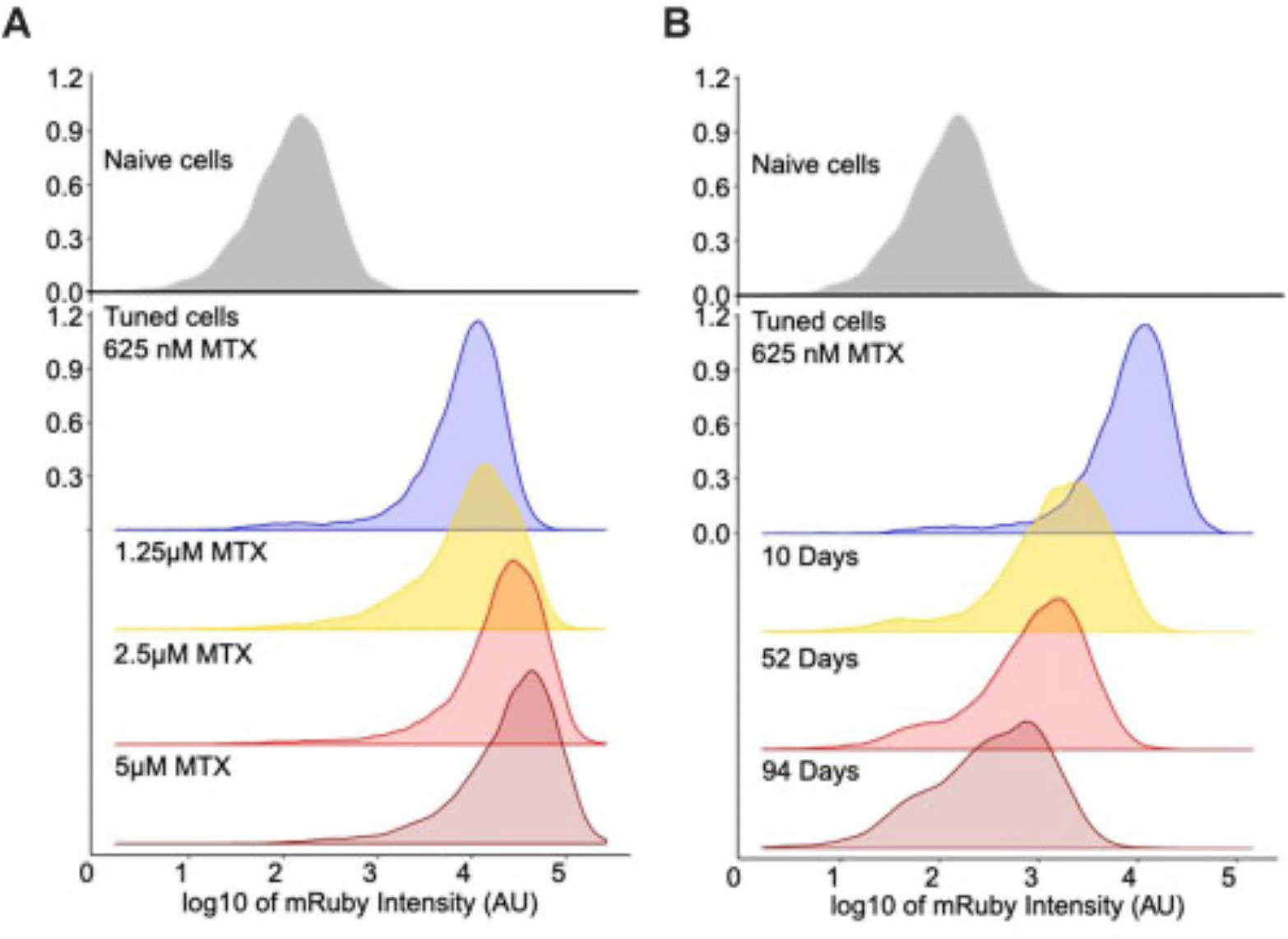
Expression of mDHFR-mRuby is commensurate to the concentration of MTX and cells revert towards the naïve state upon MTX removal. (**A)** Flow cytometry analysis shows a substantial increase in mDHFR-mRuby expression in tuned cells under 625 nM MTX challenge. Moving these tuned cells to higher concentrations of MTX leads to commensurate increases in mDHFR-mRuby expression. **(B)** Moving cells that have tuned to 625 nM MTX back to non-selective media shows gradually declining mDHFR-mRuby expression towards the naïve state.

Our observations are consistent with stochastic tuning of mDHFR-mRuby expression in response to MTX in the absence of a pre-existing regulatory program. However, an alternative mechanism could be accumulation of mutations in the local promoter sequences. By sequencing five colonies, we showed that the observed tuning is not due to any mutations in the regulatory region. In addition, we used qPCR to compare the genomic copy number of mDHFR-mRuby from the tuned cells relative to naïve cells. In five independently tuned colonies isolated, only one colony had 2-3 fold increase in its copy number which, by itself, cannot account for the extreme increase in the expression of mDHFR-mRuby.

As predicted by our stochastic tuning model in yeast, tuned cells eventually revert back to the naïve state following removal of the challenge^18^. To check for evidence of this reversion in mammalian cells, we moved the tuned cells to a media without MTX. If any genetic mutations were responsible for increasing mDHFR expression, the phenotype should be stable upon MTX removal for many generations. As can be seen in **Figure 3B**, the mRuby signal diminishes over time. To test whether the lower mRuby state is similar to the naïve state, we sorted the cells with the lowest 10% of mRuby fluorescence in the reverting population by using Fluorescence activated cell sorting (FACS). Most of the cells in this 10% low end of mRuby fluorescence died after re-exposure to MTX and roughly ∼1% tune again over a period of two weeks, consistent with a non-mutational process. In addition, we followed a small number of tuned cells, sorted to be in the top 10% of mRuby expression, in the media without MTX over a span of five days using fluorescence microscopy. These cells showed declining fluorescence over time in the absence of MTX. This provides further evidence that the tuning is reversible and that the decreased fluorescence (**Figure 3B**) is not due to a simple selection process, as it occurs at the level of individual cells over a period of days. The reversion to the naïve state, and the subsequent ability to tune again provides a strong indication that the increase in the expression of mDHFR-mRuby is due to a non-mutational adaptation process consistent with stochastic tuning.

## Discussion

Regulation of gene expression is fundamental to all cellular processes—from stress response in bacteria, to precise orchestration of gene expression during development. As such, much of the genome encodes sensory, signal transduction and regulatory functions, forming complex gene regulatory networks (GRNs) that activate the expression of encoded biochemical functions to match the requirements of a constantly changing environment. Such hard-wired GRNs are thought to arise over long-term evolution when expression of a gene product confers significant fitness advantage under a specific frequently encountered environmental context^29,30^. However, how does an organism adapt to challenges it has rarely (or never) experienced in its evolutionary history? We previously proposed that cells could adapt to such scenarios by using noise to randomly increase or decrease the expression of individual genes and to actively reinforce those changes that improve the overall health of the cell^18^. We provided evidence that yeast cells indeed utilize this novel form of adaptation we have termed stochastic tuning^18^. Here, we explored the possibility that mammalian cells also utilize stochastic tuning to adapt to unfamiliar/extreme challenges beyond the capacity of their genetically encoded pathways. We developed a simple system that makes survival/growth of human HEK293 cells dependent on the expression of a methotrexate-resistant mDHFR gene driven from a weak synthetic promoter. We observed that when these cells are exposed to the antimetabolite drug methotrexate, they manifest an adaptation process entirely consistent with stochastic tuning.

Key findings from our study include the ability of cells to tune mDHFR-mRuby expression levels in direct proportion to the concentration of MTX, enabling them to survive and grow in the presence of otherwise lethal concentrations of MTX (**Figures 2 and 3**). Consistent with our observations in yeast, only a fraction of cells successfully tune and grow in the presence of otherwise lethal MTX levels. This is consistent with the non-deterministic nature of a trial-and-error process in which success declines with the severity of challenge imposed^18^. Importantly, we show that the dramatic increase in expression of mDHFR-mRuby is not driven by promoter mutations or significant copy number variation, supporting a non-mutational mechanism consistent with stochastic tuning. Upon removal of MTX, cells reverted to their naïve state, as indicated by diminished mRuby fluorescence. This reversion underscores the dynamic nature of stochastic tuning, distinguishing it from permanent genetic alterations. Moreover, the re-tuning capability observed in reverted cells highlights the potential for stochastic tuning to provide a flexible mechanism for navigating fluctuating environments. Our observations of gradual increases in mDHFR-mRuby expression and the ability of individual cells to tune and re-tune their expression argue against a purely selection-based mechanism. Instead, our findings are more consistent with stochastic tuning, a process by which random transcriptional fluctuations are reinforced to improve the overall health of the cell.

Our findings have significant implications for understanding non-mutational mechanisms of chemotherapy resistance. The ability of cells to dynamically upregulate resistance-inducing genes, such as mDHFR, in response to drug exposure suggests that stochastic tuning may contribute to the emergence of drug-resistant populations under exposure to other cytotoxic or targeted therapies. Here, we have developed a contrived system of stochastic tuning by challenging cells to express a resistant variant of mDHFR from a weak synthetic promoter. However, in the clinical setting, variation in the expression of many genes (e.g. efflux pumps, drug targets, and DNA repair pathways) could contribute to overall drug resistance in a more distributed fashion. Unlike genetic mutations, which are often irreversible, stochastic tuning allows for transient and reversible adaptations, potentially providing cancer cells with a survival advantage in the face of dynamic therapeutic challenges. By enabling dynamic and reversible adjustments to gene expression, stochastic tuning complements GRNs and provides a powerful strategy for navigating extreme and unfamiliar environments. Our findings open new avenues for investigating the role of stochastic tuning in other multicellular contexts, including physiological adaptation, developmental dynamics, cancer initiation, and aging. A high priority is elucidating the detailed molecular mechanisms of stochastic tuning, the targeting of which may provide significant therapeutic benefit for cancer patients.

## Acknowledgements

We thank the Tavazoie laboratory for technical support and enlightening discussions throughout the project. S.T. was supported by NIH/NIGMS grant R01GM139215. A.M.R was supported by a Ruth Kirschstein NRSA Postdoctoral Fellowship Award (F32-GM125170).

## Methods

### Constructs

We first generated a minimal expression reporter by replacing the eGFP cassette in pBI-MCS-eGFP^31^ (Addgene #16542) with an mDHFR(L22R)-mRuby fusion. The L22R murine DHFR variant is resistant to methotrexate. The minimal CMV promoter used here is a truncated version of the CMV core promoter which exhibits minimal expression in HEK293 cells^32^. To generate the reporter construct, we excised the CMV promoter and ORF region of a pCDNA5/FRT vector (Thermo Fisher) and replaced it with the tet-off-minimal CMV-mDHFR-mRuby cassette cloned from the pBI based construct. We refer to this construct as the pcDNA5/FRT reporter. As a negative control, an inactive mDHFR-mRuby variant was created by deleting amino acids 20–34 of mDHFR.

### Stable Cell Line Construction

To generate the reporter line, the pcDNA5/FRT reporter construct was cloned into Flp-In 293 (Life Technologies), a human cell line related to human embryonic kidney 293 cells that contains a single integrated FRT site at a transcriptionally active genomic locus. Flp-In 293 cells were co-transfected with the pCDNA5/FRT-based plasmid and plasmid pOG44 (Life Technologies), which expresses the Flp-In recombinase. Transfections were performed using Lipofectamine 2000 according to the manufacturer’s recommendations (Life Technologies). Stable cell lines expressing the reporter system were selected for 10–12 days under 50 μg/ml of hygromycin. To generate a stable cell line expressing tTA, the tTA-IRES-Neo plasmid (Addgene plasmid #16541) was linearized using BglII and transfected into Flp-In 293 cells using Lipofectamine 2000, and a stable cell line was selected using 800 μg/ml of G418.

### Preparation of Methotrexate Solution

Methotrexate di-sodium salt (250 mg, Alfa Aesar) was dissolved in 10 mL of water to prepare a 0.5 M stock solution.

### Stochastic Tuning Assay

For stochastic tuning trials, lightly seeded 293 cells were cultured for one day in 2 mL of DMEM (Thermo Fisher) in six-well plates. Methotrexate solution was added to the wells on the second day at the desired concentrations. The culture medium was refreshed every 3–4 days with fresh DMEM containing methotrexate. Individual colonies were picked using sterile TrypLE-saturated filter paper (Thermo Fisher) and transferred to fresh media for expansion.

### Fluorescence-Activated Cell Sorting and Analysis

Cells were harvested at 90%–95% confluency for sorting and analysis. Fluorescence-activated cell sorting (FACS) was performed using a BD FACSaria sorter with the mRuby fluorescence channel for detection.

### qPCR Analysis

To measure the fold change of the mDHFR cassette copy number, DNA was extracted from HEK293 cells using the Qiagen Blood & Cell Culture DNA Mini Kit. The primers used for qPCR amplification of the mDHFR cassette were CCTGGTTCTCCATTCCTGAG and TTCCTGCATGATCCTTGTCA. qRT-PCR was performed using SYBR® Green PCR Master Mix (Life Technologies, Carlsbad, CA, USA) and the QuantStudio 5 system. The ΔΔCt method was used to determine the relative copy number of the target gene compared to the ACTB housekeeping gene using primers AGAGCTACGAGCTGCCTGAC and CGTGGATGCCACAGGACT.

